# Stuck in the cell: trapping viruses in infected cells was a frequent adaptation strategy in human hosts

**DOI:** 10.64898/2025.12.12.693836

**Authors:** Mary Reed-Weston, Jesús Murga-Moreno, David Enard

## Abstract

Hosts and viruses are in a constant evolutionary arms race, in which viruses physically interact with many host proteins (immune and non-immune) in order to replicate. Here, we manually classify ∼2,500 human virus interacting proteins (VIPs) as a function of their involvement during specific viral replication steps in order to quantify host adaptation across the viral replication cycle. We use an extension of the McDonald-Kreitman test, ABC-MK, to quantify adaptation in human VIPs compared to confounder-matched non-VIPs at different steps of the viral replication cycle. We find significant adaptation at viral replication cycle steps related to entry and release of virions from the cell, with release in particular having experienced extremely high adaptation during human evolution. In contrast to the strategy of host adaptation at entry, which prevents the virus from getting in the cell and replicating in the first place, adaptation at the release stage traps new virions in the cell to prevent continued infection. We explain this unique pattern of adaptation under the framework of a process called intergenerational phenotypic mixing. This adaptation that prevents new viral particles from exiting the cell to continue infection in the host was such a frequent and important strategy in human hosts that the evidence of it can still be found in the form of extremely strong measures of positive selection.

**Author Summary:** Viruses have been a powerful driver of evolution throughout human history. Though there have been many studies that have examined the interactions between viruses and the human immune system, few have approached the question from the perspective of what the virus needs from a host cell in order to replicate. As a virus interacts with many host proteins of various functions, whether it is hijacking or suppressing the normal cellular functions of these proteins, identifying adaptation where human hosts defeated viruses in the past may provide key insights that could lead to a better understanding of host-virus interactions, as well as new potential antiviral drug targets. Here, we test for positive selection in manually curated gene sets that span the viral replication cycle in host cells (e.g. entry into the cell, replication, transport within the cell). We identify specifically the step of viral release from the cell, wherein newly assembled viral particles exit an infected cell to go infect another, as a hotspot of extremely elevated adaptation. This result gives new insight into how human hosts defended against viruses in the past and identifies specific functional classes of proteins that were the targets of repeated, strong adaptation.

## Introduction

Hosts and pathogens are in a constant evolutionary arms race, the evidence of which can be detected in the form of adaptation in host genomes [1]. Previous research on host virus-driven adaptation has primarily focused on the interactions of viruses and the individual cells that they infect from the perspective of host immune genes that combat viral infections; i.e. the host antiviral immune response [2–4]. However, viruses also interact with many non-immune host genes in order to replicate. Thus, host adaptation in response to viruses can also happen at steps of the viral replication cycle that do not involve host immune genes [1]. This includes, for example, when a virus enters a host cell, when the genome is replicated, when new virions are assembled, and when the new virions are released from the cell. The idea of viruses being controlled by host adaptation at specific steps has thus far mainly been explored at the step of receptor attachment and overall viral entry into the cell, but there may be significant adaptation at other steps of the viral replication cycle that has remained overlooked.

To date, no studies have systematically examined virus-driven host adaptive evolution from the perspective of all the generic steps that viruses go through to complete their entire replication cycle in infected host cells (Figure 1).

**Figure 1.**
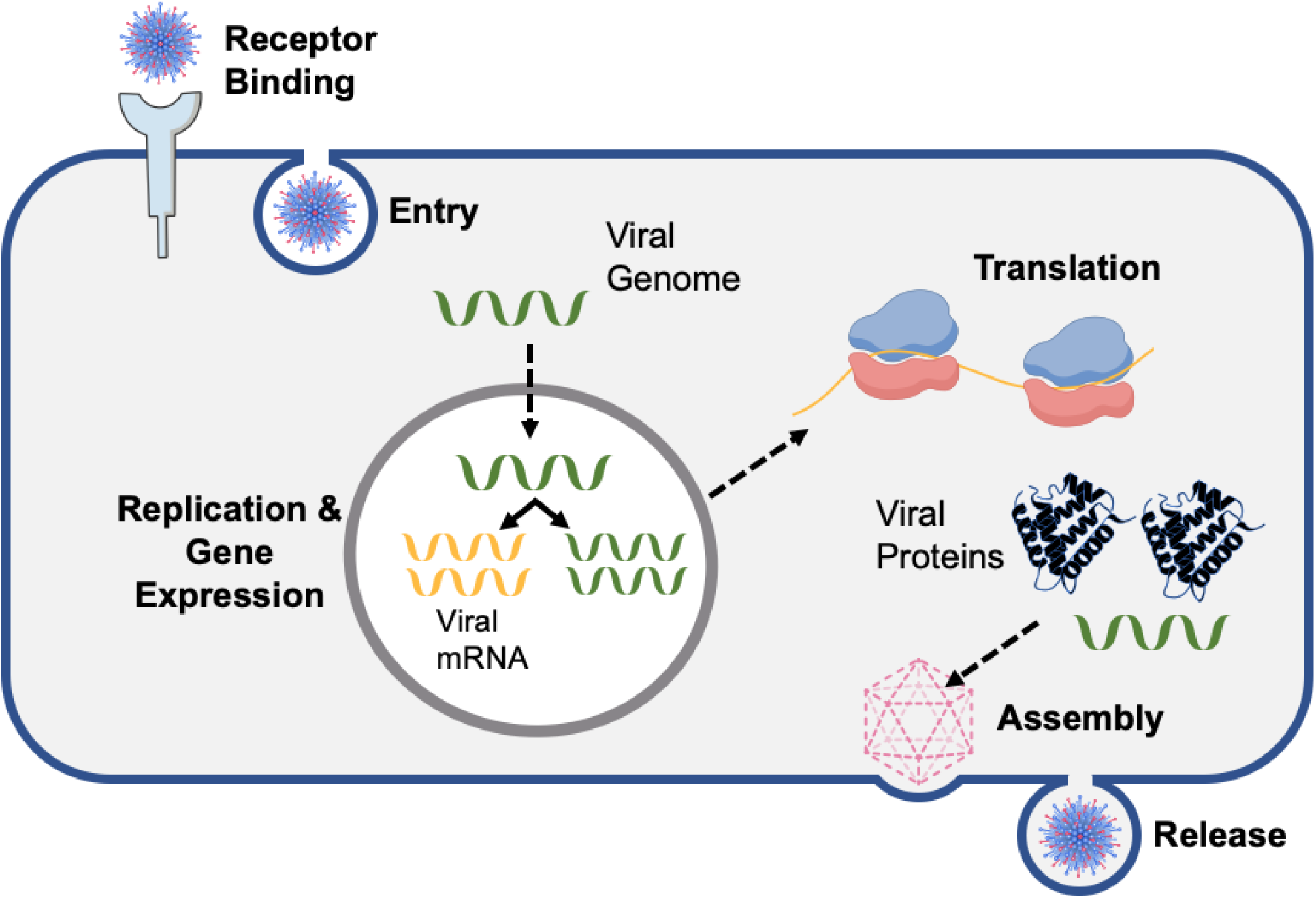
Schematic representing general viral replication cycle steps. The virus represented here is a simplistic version of influenza A. Dashed arrows indicate intracellular transport to different regions of the cell.

As the virus moves through a host cell, it physically interacts with many host proteins to suppress, subvert, or divert their functions in order to complete steps of its replication cycle. Because these physical interactions are in fact the main molecular mechanism used by viruses to subvert host cells or by the host to attack viruses, they also are the sites of a large amount of host adaptation during arms races [1, 5, 6].

These host proteins, known as virus interacting proteins (VIPs), can be examined for evidence of adaptive evolution at different steps of the viral replication cycle. To ensure that the adaptive evolution being detected and quantified is due to viruses specifically and not another selective pressure, proteins that are not known to interact with viruses (non-VIPs) can be used as a tool of comparison with VIPs.

While it has become clear that non-immune VIPs can be under strong selection during viral infection [1, 7], most well-studied instances are restricted to receptors for viral attachment. For example, the ACE2 receptor has been studied extensively in recent years due to its role relating to coronavirus (particularly SARS-CoV-2) binding and entry into the cell [8–11]. Studies have found that variants in ACE2 influence susceptibility to SARS-CoV-2 infection by reducing the binding affinity of the virus’ spike protein [12–14]. In turn, some new variants of SARS-CoV-2 in circulation have mutations in the spike protein that binds to ACE2 which increase the binding affinity and thus susceptibility to infection [15–17]. Another case of selection in a receptor protein is the NPC1 receptor, which enables filovirus attachment to host cells. One study found that, despite significant functional constraint in the NPC1 protein, there was evidence of positive selection in all mammalian orders [18]. These positively selected mutations primarily occurred within the region of the protein that binds the glycoprotein of the virus. Similarly to the case of ACE2 and SARS-CoV-2, the viruses using NPC1 also showed positive selection specifically in the protein that binds to NPC1, thereby increasing entry of the virus into the host cell [19]. In these cases, there is intense selective pressure on a non-immune VIP. However, despite these well-known case, there has been limited research into host proteins at other replication cycle steps.

The major reason why the distribution of host adaptation has not been systematically studied across viral replication steps is that there are currently thousands of known VIPs [20], each of which could be involved in one or multiple viral replication steps. This large number has previously prevented systematic efforts to comprehensively annotate VIPs; there is no manually curated database of human VIPs that classify them according to their specific role in the viral replication cycle. Here we create such a database, after the painstaking manual curation of molecular virology data over a period of ∼5 years. We then use the resulting dataset of 2,477 VIPs with at least one antiviral or proviral function to conduct a comprehensive examination of adaptive host evolution across the viral replication cycle.

In this paper, we manually classify 2,477 human VIPs as a function of their involvement during specific viral replication steps in order to quantify host adaptation across the viral replication cycle (Materials and Methods). Using categories that virologists themselves use to characterize the viral replication cycle (Figure 1), we manually browse the existing virology literature (Materials and Methods) to classify each gene encoding each VIP according to (i) replication cycle step, (ii) whether it is related to immune function or not, and (iii) whether it has a proviral (protein the virus hijacks for its own use) or antiviral (protein the virus must suppress or evade) function. These categories include: receptor binding, viral entry into the cell, intracellular viral transport, viral genome replication, viral gene expression, viral assembly, viral release from the cell, immune function, and control over the cell cycle and survival of the cell. Of 5,500 VIPs, 2,477 have identified replication cycle functions, with numbers in different categories ranging from 117 genes for viral genome assembly to 679 genes for intracellular viral transport (Table 1). Additionally, we identified 914 immune VIPs and 1,568 nonimmune VIPs, as well as 1,294 genes with a solely proviral function and 682 with a solely antiviral function. Notably, there is overlap between replication steps, because the same VIP can play a role in different replication steps in the same or different viruses, or could be used as proviral or antiviral depending on the virus. In this case, the same VIP is classified with multiple functions. Different viruses have different numbers of annotated VIPs that reflect the virology research effort dedicated to these different viruses. For example, unsurprisingly, after decades of research, HIV has many more VIPs with annotated viral replication steps than SARS-CoV-2.

**Table 1.** VIP replication cycle categories and the number of genes in each set. Testable VIPs indicate the number that are matched for confounding factors with non-VIP controls.

| Category | All VIPs | Non-immune testable VIPs |
| --- | --- | --- |
| Receptor | 128 | 67 |
| Entry (excluding receptor) | 203 | 140 |
| Viral genome replication | 334 | 227 |
| Viral gene expression | 444 | 289 |
| Intermediate transport | 138 | 97 |
| Assembly | 117 | 75 |
| Release | 261 | 154 |
| Release (excluding assembly) | 224 | 134 |
| Cell survival/proliferation | 228 | 186 |
| Immune | 914 | 790 |
| Non-immune | 1568 | 1368 |

Using these categories, we then quantify human protein adaptation since divergence from chimpanzees. To do this, we use an extension of the McDonald-Kreitman test [21]. The original version of the test compares divergence between two closely related species to the amount of polymorphism within a species at synonymous (DS: synonymous fixed differences, PS: synonymous polymorphism) and nonsynonymous sites (DN: nonsynonymous fixed differences, PN: synonymous fixed differences). The test identifies an increase in the number of nonsynonymous fixed differences (DN) due to positive selection. In order to account for variable mutation rate, DN is contrasted with the number of synonymous fixed differences (DS). The excess of DN is then estimated by comparing the DN/DS ratio to the ratio of nonsynonymous and synonymous polymorphism (PN/PS). The PN/PS ratio provides the baseline for purifying selection exceeded by the DN/DS ratio when adaptation increases DN. Because PN/PS tends to be much less than 1, this makes the MK test far more powerful to detect positive selection than tests based only on the classic dN/dS ratio, also known as the ω ratio [22, 23] that has to exceed 1 to provide evidence of adaptation. These classic ω tests would not properly quantify host adaptation across different viral replication steps, because all the adaptation that did not elevate ω above 1 would be completely neglected.

However, because the MK test uses polymorphism (PN and PS), it suffers from a number of confounding factors that do not affect classic dN/dS tests. In particular, the MK test is susceptible to the presence of weakly deleterious variants affecting PN. Using PN/PS as the baseline for purifying selection makes the assumption that PN only reflects the neutral variants remaining after all deleterious variants were removed by purifying selection. However, weakly deleterious variants can stay at non-negligible frequencies in populations for generations due to genetic drift before being ultimately removed by purifying selection. This tends to inflate PN, and make the MK test overly conservative when not taken into account. Demographic changes can also affect both PN and PS [24, 25]. To address these confounders, we use an advanced version of the MK test called ABC-MK [6, 26]. ABC-MK applies Approximate Bayesian Computation to estimate the proportion of those amino acid substitutions (DN) that were due to adaptation. This proportion is denoted α, which can be measured with the following equation (Equation 1).

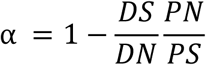

A further advantage of using the MK test for this study is that it accounts for potentially varying levels of selective constraint between the different viral replication cycle steps with the aforementioned PN/PS baseline. ABC-MK also provides separate estimates of strong amino acid adaptation that went to fixation too fast to contribute to nonsynonymous variation, and weak adaptation that was slow enough to still contribute to nonsynonymous variation before fixation [6, 25]

Along with selective constraint, the adaptive potential of genes may also depend on factors other than interactions with viruses that could confound our analysis, such as gene expression, the number of human protein interactions with other human proteins (rather than viruses), whether the gene is a housekeeping gene or not, etc (see Materials and Methods for the full list of confounding factors used). To account for this, we contrast VIPs with control non-VIPs that match the VIPs for multiple possible confounding factors that might also affect the occurrence of adaptive evolution (Materials and Methods). The difference between VIPs and control non-VIPs can then be attributed to interactions with viruses.

We use ABC-MK to quantify adaptation in human VIPs compared to confounder-matched non-VIPs at different steps of the viral replication cycle. We find significant adaptation at viral replication cycle steps related to entry, viral transport, and release of virions from the cell, with release in particular having experienced extremely high adaptation during human evolution. Though positive selection at entry and receptor proteins has been well-documented (ACE2 and SARS-CoV-2, NPC1 and filoviruses), intense selection at exit from the cell had not been identified beyond the cases of the specialized antiviral proteins tetherin and viperin. Tetherin is expressed at the surface of human plasma cells, bone marrow cells, and B cells, where it was recorded to inhibit the release of HIV-1 from the host cell [27–29]. Viperin is an interferon stimulated gene that is strongly induced in host cells in response to viral infection. Viperin has been identified as having a role in inhibiting the release of influenza A viral particles from an infected cell by disrupting lipid rafts used by the virus [30, 31]. Here, we have identified high adaptation across many proteins related to viral release, more than the previously known cases of viperin and tetherin. In contrast to the strategy of host adaptation at entry, which prevents the virus from getting in the cell and replicating in the first place, adaptation at the release stage traps new virions in the cell to prevent continued infection, as evidenced by the cases of viperin and tetherin.

## Results

We use ABC-MK to estimate the amount of protein adaptive evolution across 11 steps of the viral replication cycle. Table 1 summarizes the numbers of VIPs used to represent each step. To control for confounding factors, we use a bootstrapping method described in [32], [33], and [37] to build control sets of non-VIPs that match the lists of VIPs used for each viral replication step (Materials and Methods). In total we match 16 different confounding factors. For each set of VIPs, we generate 1,000 control sets of matched non-VIPs allowing us to estimate a null distribution of expected adaptive evolution in the absence of interactions with viruses, as well as corresponding empirical p-values (Materials and Methods). Fig. 4 shows these null distributions. However, because we match multiple confounding factors between VIPs and non-VIPs, the number of the latter that can be used as controls is significantly decreased. Such smaller sets of non-VIPs that can be used to build control sets increase the variance of the expected null distributions and therefore the risk of both type I and type II errors in our test, as previously described in [5], [32] and [34]. To account for this increased risks, we perform a simple random shuffling between VIPs and matched control non-VIPs to determine an unbiased false positive rate (FPR) of our test for each tested set of VIPs (Materials and Methods; S1 Table).

Because immune VIPs include many prominent cases of well-studied adaptation in response to viruses, we first compare the total number of adaptive substitutions in immune VIPs vs. non-immune VIPs. We surprisingly find that immune categories show minimal to no adaptation relative to the to the non-immune sets (Figure 2), a result that is nevertheless reminiscent of previous results using a more basic version of the MK test [1]. We hypothesize that there might be significant balancing selection maintaining nonsynonymous variants in the immune genes studied [1, 35]. The MK test in general and ABC-MK are expected to be biased downward in the presence of balancing selection, because the increased amount of nonsynonymous polymorphism being maintained is simply registered as an elevated PN, thus bringing down the value of α. Because of this, for the rest of our analysis we use only non-immune genes and exclude genes annotated with immune functions (Materials and Methods).

**Figure 2.**
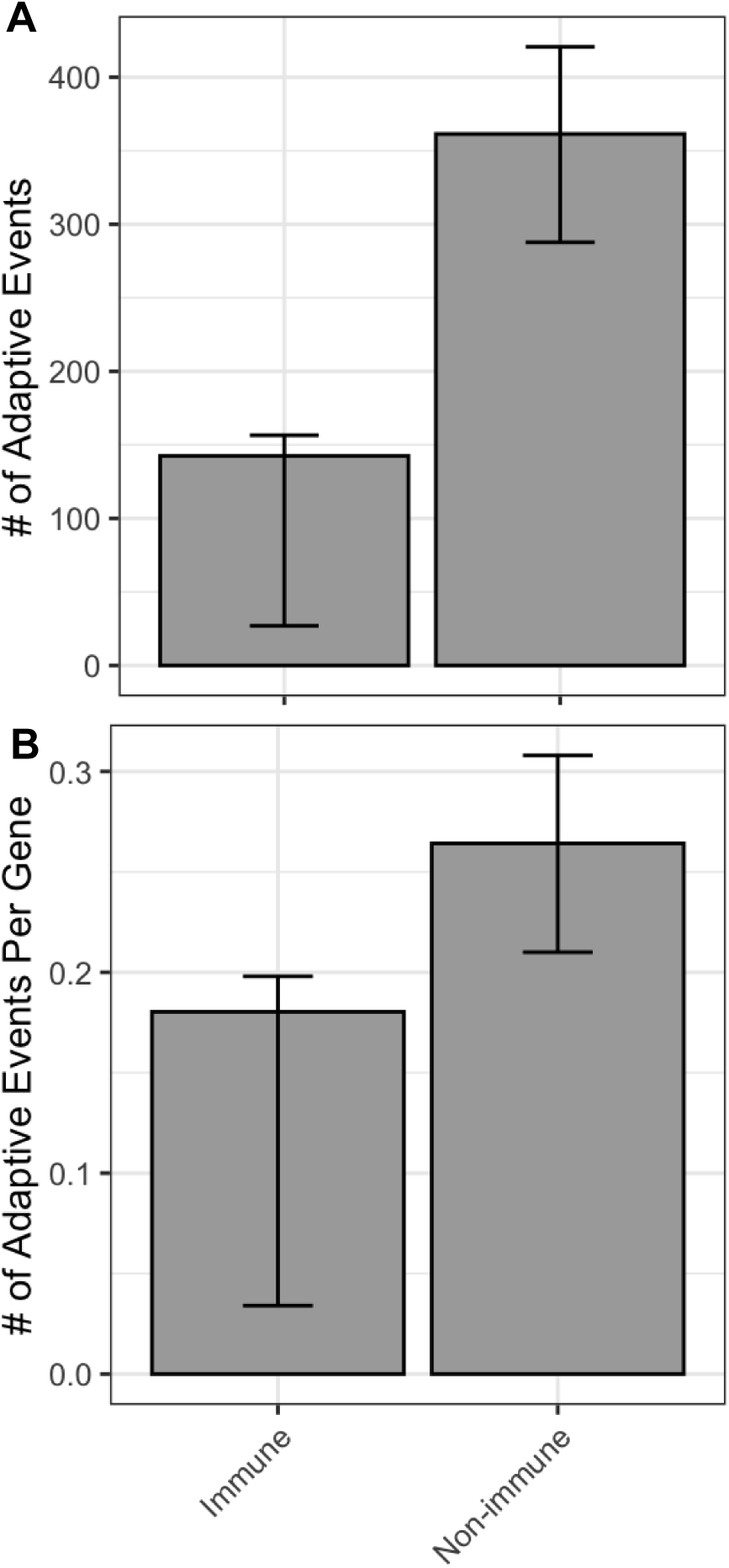
Comparisons of the number of adaptive events between genes that have an immune function and those that do not. **A:** The total number of adaptive events. **B:** The number of adaptive events per gene. Error bars indicate 95% confidence interval from resampling

Because we have not previously studied the impact of this specific potential confounding factor, we then focus on the possible impact of the housekeeping gene status [36] of VIPs compared to non-VIPs. Focus on housekeeping genes in VIPs vs. non-VIPs is justified by (i) the fact that we observe a much higher proportion of them among VIPs (43%) vs. non-VIPs (17%), and (ii) the fact that we expect *a priori* that housekeeping genes should have reduced adaptive potential. In line with this, we find that housekeeping VIPs do not show increased adaptation compared to control non-VIPs (Figure 3). In contrast, non-housekeeping VIPs have experienced increased adaptation compared to non-housekeeping non-VIPs (Figure 3). Because of this profound effect of housekeeping gene status, we add their proportion to the list of confounding factors to match between VIPs and non-VIPs (Materials and Methods). The lack of increased adaptation in housekeeping VIPs suggests that even viruses did not represent a strong enough pressure to drive adaptation in these highly constrained genes. Accordingly, we observe not only a lack of overall increased adaptation, but specifically a lack of strong adaptation (using the ability of ABC-MK to distinguish between weak and strong adaptation; Fig. 3B), which is usually the hallmark of VIP adaptive evolution [6, 37].

**Figure 3.**
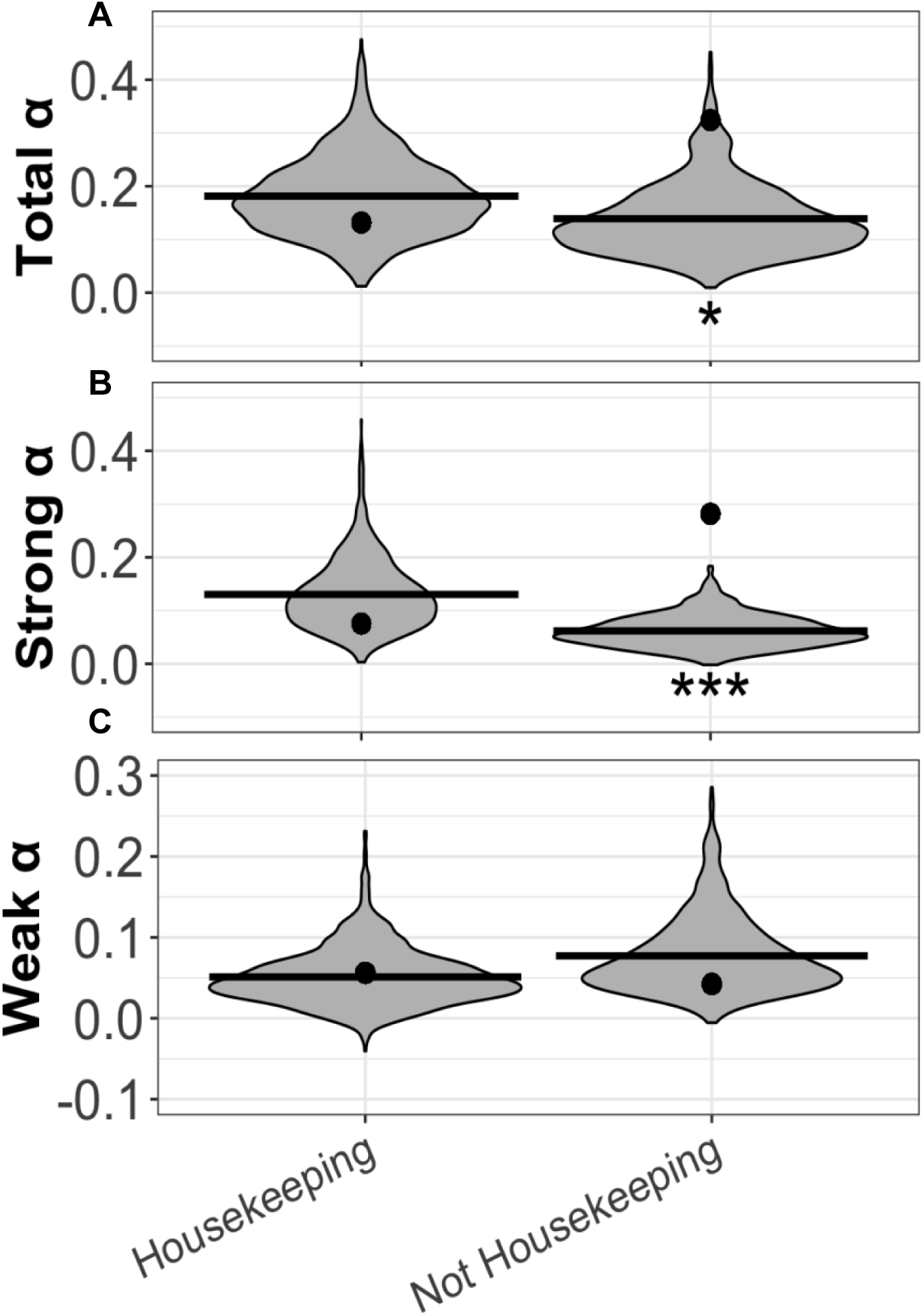
Comparisons of α values from ABC-MK between sets are entirely comprised of housekeeping genes and sets that entirely exclude housekeeping genes. Violin plots represent α values for the 1000 control sets, with the black point representing the α value for the VIP set. *: p<0.05, **: p<0.01 ***: p<0.001. **A.** Results for the total value of α. **B.** Results for strong adaptation only. **C.** Results for weak adaptation only.

Having clarified the influence of immune and housekeeping genes, we then run ABC-MK on each replication cycle category. Out of 10 categories of VIPs vs. corresponding controls, four show a significant increase (empirical p-value P<0.05) of the proportion of total adaptive amino acid substitutions (α) relative to that of non-VIP controls. Five categories show a significant (P<0.05) increase of the proportion of strongly advantageous amino acid substitutions. The significant categories according to the empirical p-value coincide with those significant according to the unbiased FPR (FPR<0.05; Materials and Methods). These significant categories are spread across the replication cycle of the virus, rather than being concentrated at a specific stage (Figure 4). In particular, the categories of entry & receptor, entry at the exclusion of receptors, overall viral transport, and viral release are all significant (P<0.05). Thus, even though different categories overlap with partially shared VIPs, this sharing is not sufficient to even out results between categories and to hide a broad pattern that the movement of viruses in and out as well as within infected cells has been a major target of human host adaptation.

**Figure 4.**
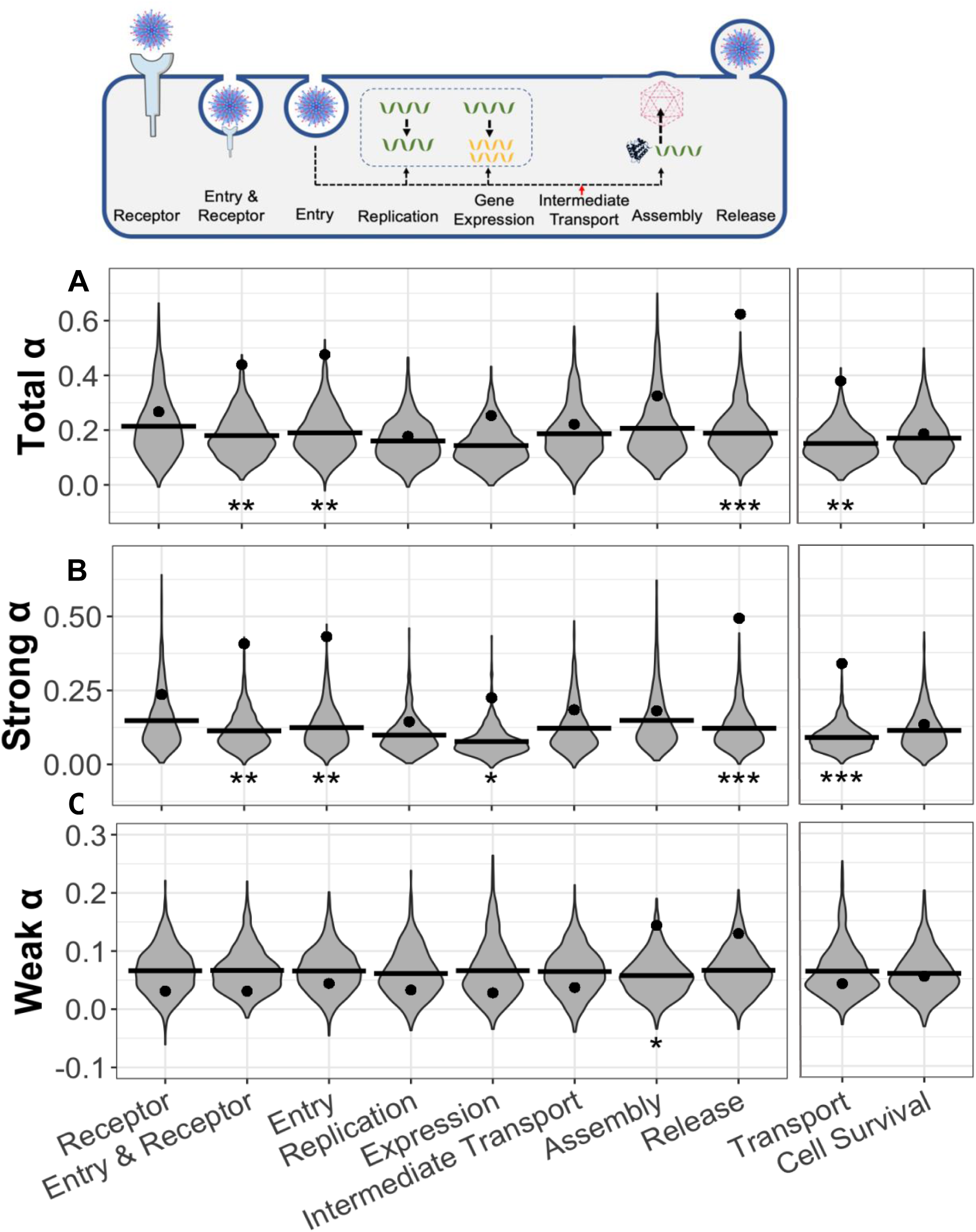
Comparisons of α values from ABC-MK for each major replication cycle category. Overlaid is a schematic of a generic virus going through its replication cycle at each of the identified replication cycle categories. Transport and Cell Survival (related to control of the cell cycle and cellular proliferation) are separated from the other categories due to being broad categories that are not represented in the schematic. Violin plots represent α values for the 1000 control sets, with the black point representing the α value for the VIP set. *: p<0.05, **: p<0.01 ***: p<0.001. **A.** Results for the total value of α. **B.** Results for strong adaptation only. **C.** Results for weak adaptation only.

The category of gene expression also shows a significant excess of strong adaptation (P<0.05). The α curve for this category supports the significant value, with the asymptote of the curve being as high as or higher than the estimate from ABC-MK (Figure S1). All significant categories show a significant excess of strong adaptation rather than weak adaptation, which we previously found to be a hallmark of host adaptation in response to viruses [6, 37].

In addition to the proportion α of advantageous substitutions, Figure 5A provides the total number of advantageous substitutions per category. Since the number of VIPs per category varies greatly (Table 1), we also provide the number of adaptive substitutions per gene in each category in Figure 5B. Of the 10 general replication cycle categories used in Figure 4, release has by far the highest number of adaptive substitutions per gene, with almost one adaptive fixation per gene (Figure 5B). The α curve for release VIPs supports this high value, with the curve for VIPs being distinctly above the curve and confidence interval for release non-VIPs (Figure 6). This is notably higher than the entry/receptor category that has previously attracted most of the attention on host adaptation. These data examining the amount of adaptation per gene correlate well with the ABC-MK results, indicating that the patterns we see in Figure 4 reflect the relative proportion differences, but also the absolute differences in adaptation between the different categories.

**Figure 5.**
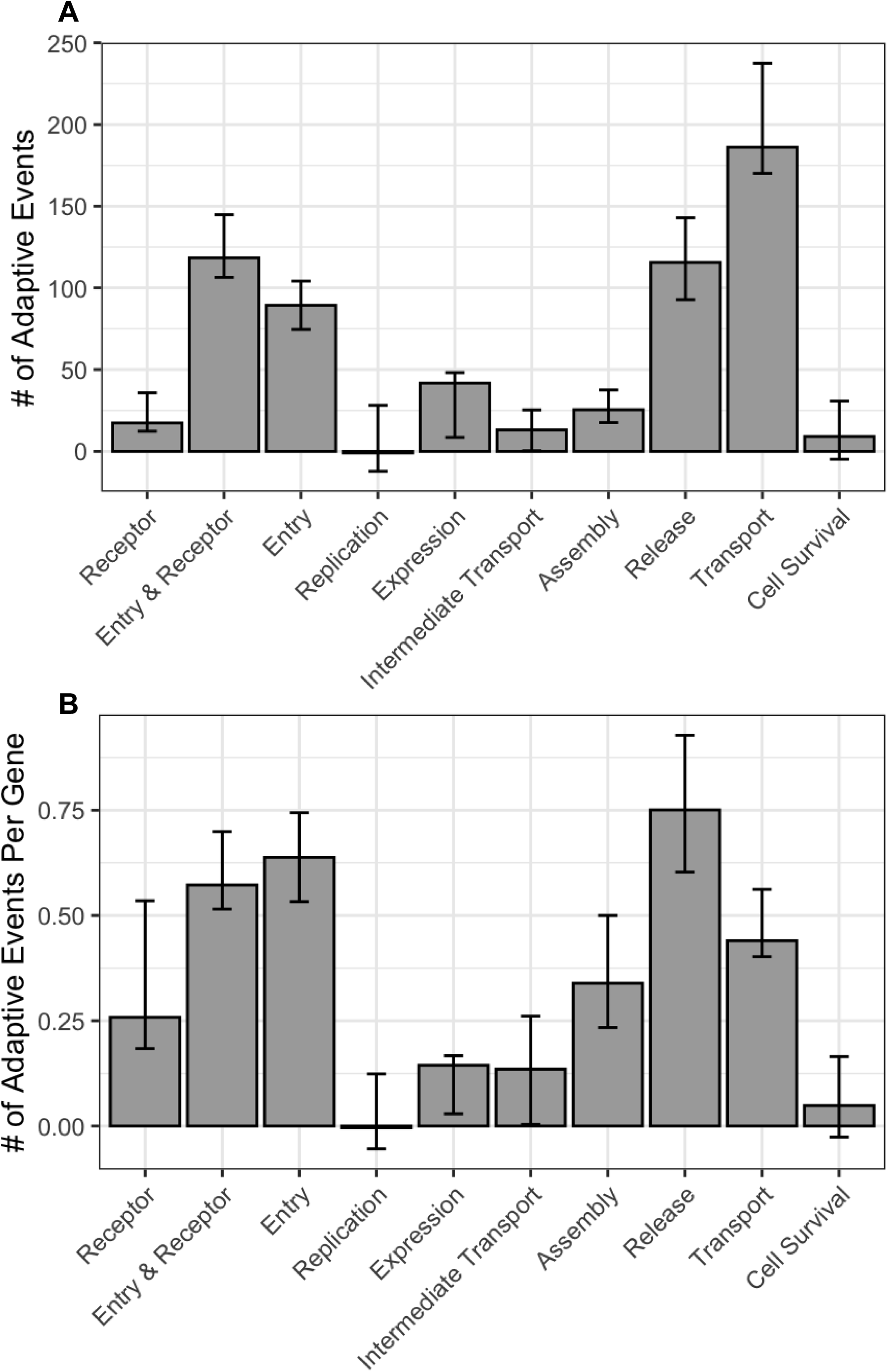
**A:** Number of adaptive fixations in each major replication cycle category. **B:** Number of adaptive fixations per gene in each major replication cycle category. Error bars represent the 95% confidence interval from resampling.

**Figure 6.**
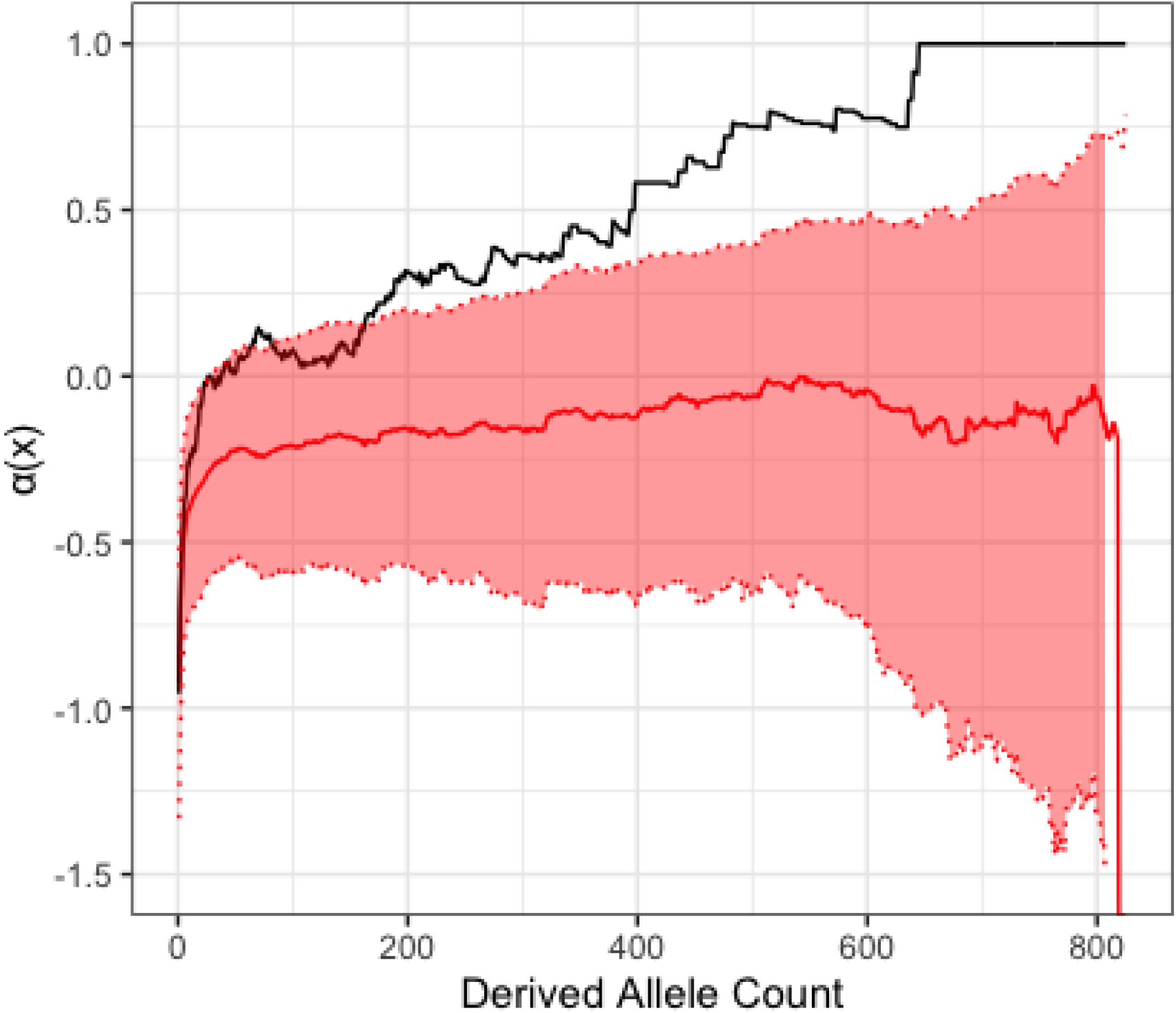
α(x) curve for release VIPs (black) compared to the mean alpha curve for release nonVIPs (red). 95% confidence interval for nonVIPs is indicated by shaded region.

## Discussion

In this study we have quantified adaptation at different steps of the viral replication cycle using human 1000GP data in the first comprehensive view of adaptation across the viral replication cycle. Our results show that we can distinguish specific steps of the viral replication cycle that prompt higher host adaptation than others even in human data, which has relatively low genetic diversity compared to other species [38, 39], and therefore lower statistical power to detect different amounts of adaptation. This is evidenced by our use of an ABC extension of the McDonald-Kreitman test, showing that categories related to viral entry and receptor binding, viral entry excluding receptor proteins, viral transport, and above all viral release have much higher values of α than their matched control sets (P<10^-3^, P<10^-3^, P< 10^-3^, P<10^-4^, respectively). While we use ABC-MK only on sets of genes and not individual genes, we can use the data provided to ABC-MK (PN, PS, DN and DS) for individual genes to identify specific genes with a DN/DS ratio greater than PN/PS, and of potential interest for future study.

In the case of viral entry, the result of significant host adaptation is in congruence with previous studies that found high host adaptation in receptor proteins that facilitate viral entry into the cell [9, 18, 40]. However, our results show that it is not just specific receptor proteins that are under strong positive selection, but rather many genes across the entry category from receptor binding all the way into the cell. For example, the VIP in the entry & receptor category (that has no other identified function) with the highest DN/DS relative to the PN/PS ratio is PLG, a protease that contributes to the infectivity of IAV. PLG cleaves the glycoprotein hemagglutinin (HA), which extends from the surface of the viral envelope of IAV. The cleavage of HA is required for IAV to infect host cells, making PLG crucial for viral infectivity and replication [41]. PLG has 6 nonsynonymous substitutions, a DN/DS ratio of 2, and a PN/PS ratio (derived allele frequency >10%) of 0 (Table 2). Table 2 provides other examples of entry genes (that have no other viral function) with elevated DN/DS compared to PN/PS.

**Table 2.** Genes of interest related to viral entry and viral release. For all genes, DN/DS is at least twice as large as PN/PS, and polymorphisms below frequency of 10% are excluded. Genes classified as either entry or release have only that function, while those classified as both have only those two functions. DN & DS refer to the number of nonsynonymous and synonymous fixed substitutions, respectively. PN & PS refer to the number of nonsynonymous and synonymous polymorphisms, respectively. EBOV: Ebola virus, IAV: Influenza A virus, RVFV: Rift Valley fever virus, HASTV: Human astrovirus, HIV: Human immunodeficiency virus, DENV: Dengue virus, HSV: Herpes simplex virus, PRV: Pseudorabies virus, LASV: Lassa virus, SEV: Sendai virus, HBV: Hepatitis B virus, PEDV: Porcine epidemic diarrhea virus

| Ensembl ID | Gene name | DN | DS | PN | PS | Function | Virus |
| --- | --- | --- | --- | --- | --- | --- | --- |
| ENSG00000111670 | GNPTAB | 4 | 4 | 0 | 2 | Entry | EBOV |
| ENSG00000122194 | PLG | 6 | 3 | 0 | 3 | Entry | IAV |
| ENSG00000123384 | LRP1 | 6 | 16 | 0 | 6 | Entry | RVFV |
| ENSG00000134243 | SORT1 | 3 | 1 | 0 | 3 | Entry | Rotavirus |
| ENSG00000155660 | PDIA4 | 2 | 5 | 0 | 2 | Entry | HASTV |
| ENSG00000213983 | AP1G2 | 2 | 3 | 0 | 1 | Entry | HIV |
| ENSG00000128833 | MYO5C | 3 | 4 | 1 | 4 | Release | DENV |
| ENSG00000139641 | ESYT1 | 4 | 4 | 0 | 1 | Release | HSV |
| ENSG00000144824 | PHLDB2 | 7 | 5 | 0 | 2 | Release | PRV |
| ENSG00000150995 | ITPR1 | 7 | 16 | 0 | 5 | Release | HIV |
| ENSG00000151929 | BAG3 | 4 | 1 | 1 | 2 | Release | EBOV, LASV |
| ENSG00000169855 | ROBO1 | 4 | 9 | 0 | 1 | Release | HIV |
| ENSG00000187391 | MAGI2 | 2 | 5 | 0 | 3 | Release | HIV, SEV |
| ENSG00000074695 | LMAN1 | 2 | 7 | 0 | 1 | Entry, Release | Arenavirus, HBV |
| ENSG00000184012 | TMPRSS2 | 7 | 1 | 2 | 3 | Entry, Release | SARS-CoV-2, PEDV |

In the case of release, the category that has the highest amount of per-gene adaptation, we identify the following genes with the highest number of nonsynonymous substitutions (DN) relative to the PN/PS ratio that are involved in viral release: (i) TMPRSS2, a gene known primarily for its role in the entry of SARS-CoV-2 into new cells [42] that also has a role in release of PEDV (Porcine Epidemic Diarrhea Virus) from the cell. TMPRSS2 not only enhances release of the virus from an infected cell into the extracellular space, but also augments the fusion of infected and uninfected cells to allow direct transmission of viral particles [43]. TMPRSS2 has 7 nonsynonymous substitutions, a DN/DS ratio of 7, and a PN/PS ratio (derived allele frequency >10%) of 0.67. The involvement of TMPRSS2 in both entry and release depending on the virus renders any interpretation of adaptation ambiguous. (ii) ITPR1, a calcium channel receptor in the endoplasmic reticulum that assists with the budding of new HIV viral particles from infected cells [44, 45]. ITPR1 has 7 nonsynonymous substitutions, a DN/DS ratio of 0.434, and a PN/PS ratio (derived allele frequency >10%) of 0 (Table 2). The high number of amino acid changes in the human lineage is particularly remarkable given that ITPR1 is very strongly conserved across mammals [46] (DN/DS =0.016 in primates, DN/DS =0.015 in rodents, DN/DS =0.021 in carnivores). Six of the seven amino acid changes occur in the main canonical, 2758 amino acid long isoform of ITPR1, all within a specific domain called ARM3, which spans 21.8% of the isoform [47]. This concentration of substitutions is highly unlikely given the uniformly high constraint across the sequence (binomial test P∼10^-4^), strongly suggesting the action of positive selection. The ARM3 domain plays an important role in the conformational changes during opening and closing of the calcium pore of ITPR1 [48].

Other than TMPRSS2, there is only one gene with DN/DS elevated over PN/PS that has solely entry and release functions (at the exclusion of other viral functions; Table 2): LMAN1. LMAN1 encodes the protein ERGIC-53, an intracellular cargo receptor that arenaviruses, coronaviruses, and filoviruses use for entry into the cell [49]. Without this protein, virions can form but are unable to attach to host cells and are thus noninfectious. ERGIC-53 also plays a role in the release of new HBV virions from the cell by interacting with a glycoprotein on the HBV envelope, allowing new virions to exit the cell [50]. LMAN1 has 2 nonsynonymous substitutions, a DN/DS ratio of 0.286, and a PN/PS ratio (derived allele frequency >10%) of 0 (Table 2).

It must also be noted that of the viral functions identified in Table 2, 16 out of 19 are associated solely with RNA viruses. Indeed, all of the identified genes associated only with viral entry are related to RNA viruses, and the two genes with both entry and release functions are associated with RNA viruses in the context of viral entry (Table 2). This lends support to previous findings that RNA viruses seem to be important drivers of adaptation in humans [26], and specifically connects this high adaptation to proteins related to viral entry into the cell. However, further research must be done in investigating the relationship between RNA viruses and their human hosts to draw more concrete conclusions.

Given our results with the category as a whole and with the specific examples above, it may seem surprising that such a striking pattern of adaptation at release proteins has not been observed before. As all VIPs are subject to higher constraint than nonVIPs [1], we generated site frequency spectra for release VIPs and all VIPs to determine if release VIPs seem to be under even higher constraint, something that could have limited previous detection of adaptation in release VIPs based solely on classic DN/DS tests (Figure 7). Compared to all VIPs, release VIPs show a rapid decrease in the number of polymorphisms relative to the number of substitutions at higher derived allele frequencies (Figure 7A, C vs B, D), a pattern that indicates strong purifying selection. Despite this increased constraint in release VIPs, ABC-MK still detects pervasive positive selection. Had we just used the DN/DS ratio to detect positive selection, the category of release, our most significant and unique result, would not have shown significant adaptation (DN/DS << 1), despite there being individual genes with high DN/DS compared to PN/PS.

**Figure 7.**
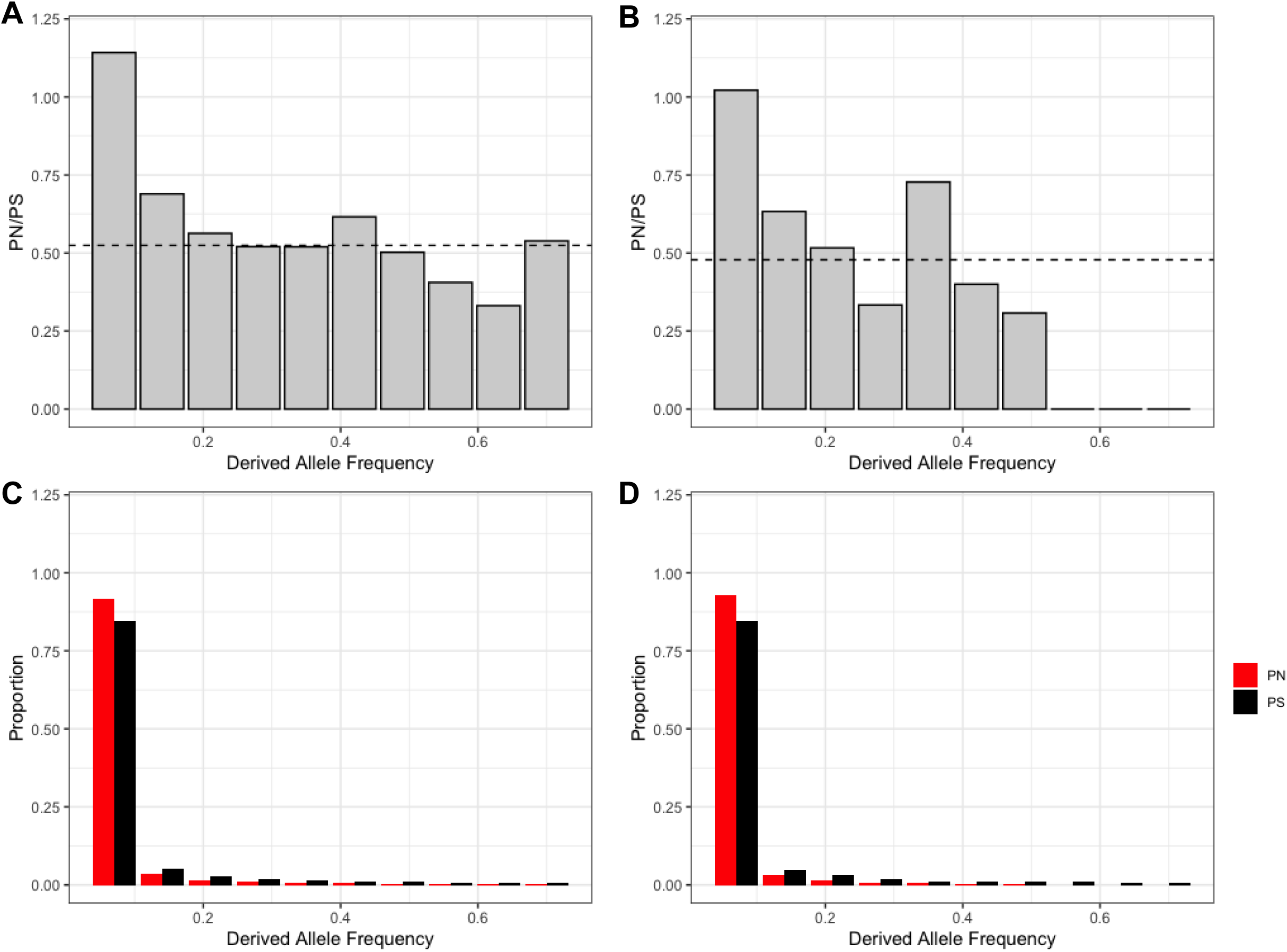
Site frequency spectra for release VIPs and all VIPs. **A-B**: Gray bars indicate the PN/PS ratio at changing variant frequencies. Black dashed lines indicate the DN/DS ratio for all VIPs (A) and release VIPs (B). **C**: Site frequency spectrum for all VIPs using synonymous (black bars) and nonsynonymous (red bars) polymorphisms. **D:** Same as C but using only release VIPs. Variants above frequency 0.7 were excluded to prevent bias by mispolarization.

The question that remains then is, why viral entry and viral release? A good adaptive host mutation is one that the virus cannot easily counteract or circumvent with its own adaptive mutations. When considering why the chronological first and last steps of the viral replication cycle (entry and release, respectively) both have high adaptation, we must consider the perspective that mutations at entry and release both are so beneficial to the host because entry is actually a *late* step, occurring after release when the new virions enter a new, uninfected cell. This perspective is significant under the framework of a process called intergenerational phenotypic mixing, viral genomes do not get assembled into virions together with the proteins that they generated, but instead with the proteins originating from other viral genomes in the same infected cell [51]. In this framework, entry into a new cell is the step furthest away from the occurrence of viral genome replication when new viral adaptive mutations can occur with the potential to counteract previous host adaptations. Release from that previous generation of cells is the second furthest step away from viral replication. We hypothesize that there is such significant host adaptation at release and entry because under intergenerational phenotypic mixing, these two steps are expected to be the ones with the greatest disconnect between the viral genomes and the viral proteins that they are packaged with. This could make adaptation more challenging for the virus, because at the stages of release and entry into a new cell, the new virion particle is already assembled and is locked into its specific combination of genome and surrounding proteins, including capsid proteins. Even if there had been an advantageous mutation that arose in a viral genome copy as it was replicated, if this new mutation is not present in enough genomes, it would likely be packaged with the old, non-adapted proteins and capsid. This delays the appearance of the new adaptive phenotype until the virion can enter a new cell, and puts the virus at a disadvantage when trying to enter the cell and replicate without the advantageous mutation(s) affecting phenotype. This may also reduce the initial genome replication of the virus in the next cell, as it cannot jumpstart replication as effectively with the old, unmutated proteins that it already has to initially express the viral genes needed.

For this reason, we expect that for a virus to overcome host adaptation at the release or entry step, the adaptive mutation must occur very early during viral replication, so that many new genomes and proteins generated from them have the mutation. This would allow the virus to overcome disadvantageous phenotypic mixing by having many mutated proteins and genomes to package into new virions. The virus needing very early mutations benefits the host because it defeats the ability of the virus to rely on very large intracellular population size as a source of new variants. In other words, it does not matter if there are many new advantageous viral variants if they are at such low frequency that they are subject to strong phenotypic mixing. What is still unknown, however, is what proportion of adapted proteins is sufficient to overcome host adaptation at release and entry, i.e. what degree of phenotypic mixing can be tolerated that still allows the virus to overcome host adaptation preventing viral release from the cell or entry into the next (Figure 8).

**Figure 8.**
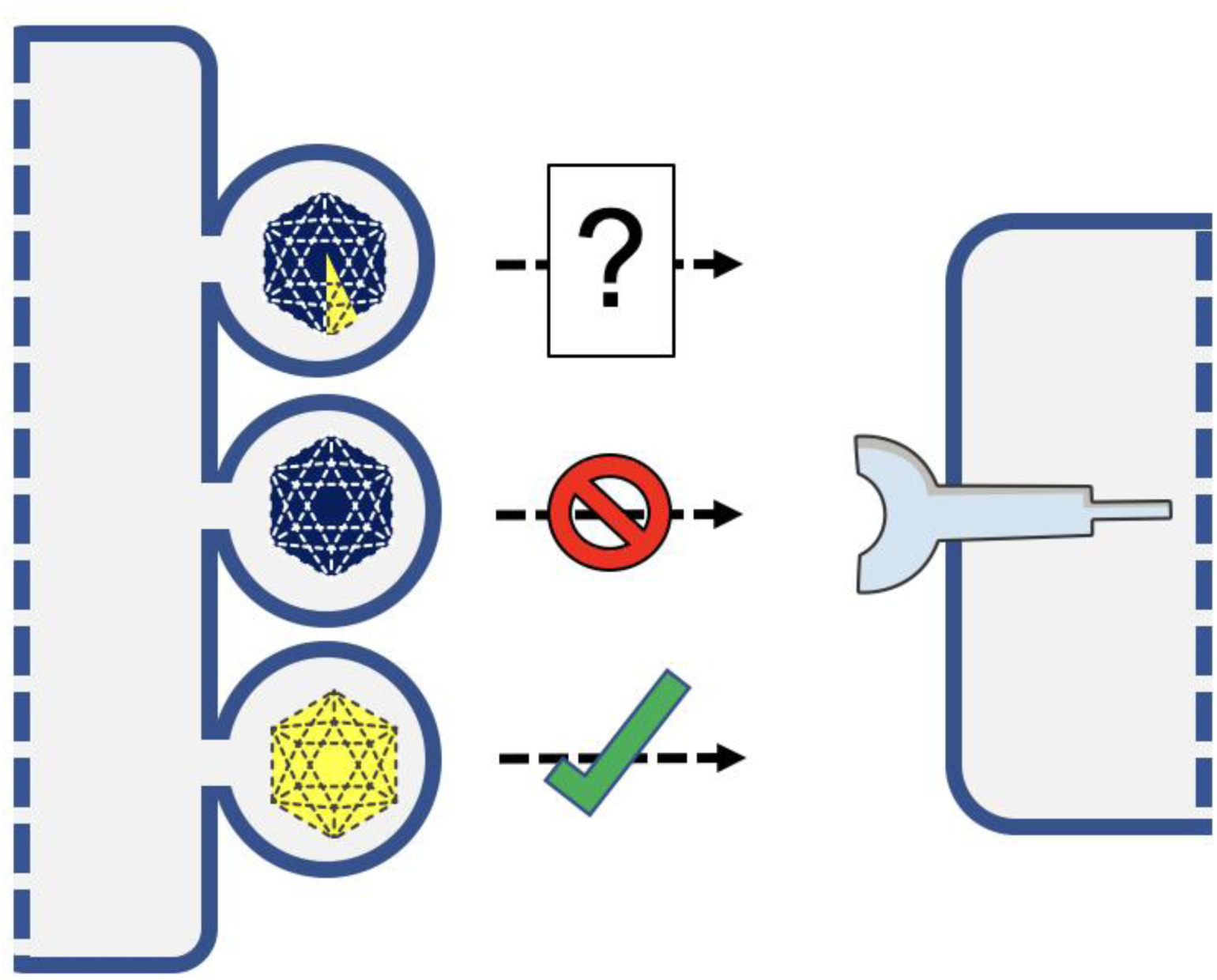
Diagram depicting the different possibilities at viral release from the cell with varying levels of phenotypic mixing. Yellow indicates viral proteins that have an adaptive mutation overcoming host adaptation at release. Blue indicates the wild-type protein. A virion with a capsid made entirely of adapted proteins will succeed in getting out of the cell and moving to the next one (likely a very early mutation). A capsid with all wild-type proteins will prevent the virus from leaving the original infected cell (no mutations or a very late mutation). The effect of a mixture of wild-type and adapted capsid proteins is unclear and requires future empirical studies.

If there is significant host adaptation at the release step, the new viral mutation might not make it out of the infected cell in the first place for the reasons described above. However, even if it does get out and proceeds to the next cell, any viral mutation that counters host adaptation at the release step may then still be more likely to be removed by host antiviral factors at all replication cycle steps it must go through again to successfully infect a new cell. As the likelihood of being eliminated by the host antiviral response likely goes up, the amount of interactions possible between different viral components at each steps decreases, namely at release and entry into the next cell.

With a mix of adapted and non-adapted viral proteins and genome copies in the cell, the virus may have multiple tries to proceed with its replication by subverting host adaptation and the immune response before the viral capsid is fully assembled and its conformation locked in place. Especially when the distribution of adapted and non-adapted proteins and genomes is not homogenous within the cell [52, 53], this allows for amplification of the mutant genotype (Figure 9). This amplification could be literal as in the case of replication, where the mutant genome is being replicated and its frequency directly increased (Figure 9A). It could also be indirect, such as in a case where mutant and wild-type proteins cannot function together, allowing the mutant proteins to be selected for. This could increase the number of mutant virions without increasing the actual number of mutant genomes or proteins, for example at the step of virion assembly (Figure 9B).

**Figure 9.**
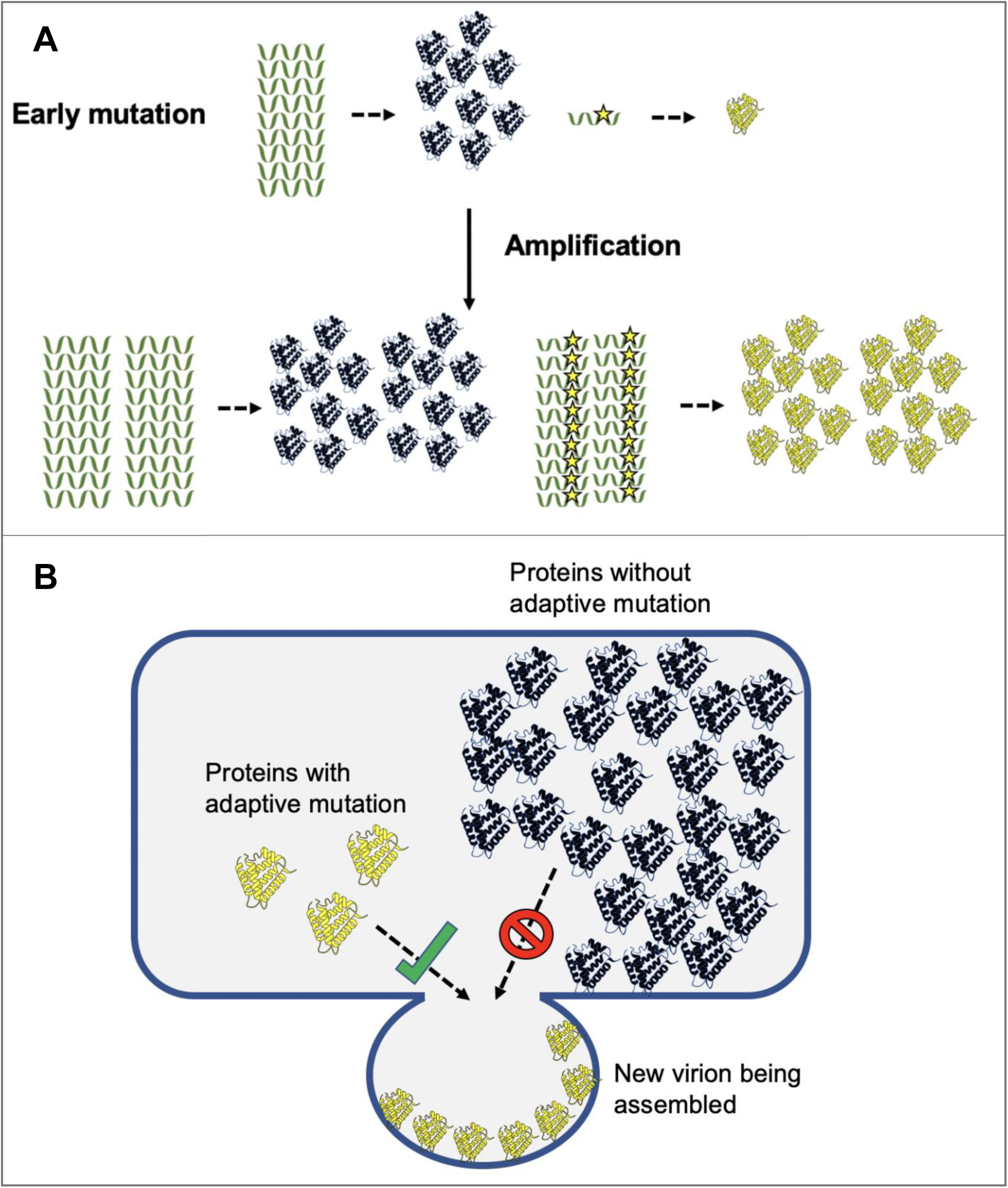
Schematics representing how different steps in the viral replication cycle can amplify a mutant viral genome. **A.** Depiction of how viral genome replication and transcription/translation increase the frequency of an adaptive viral mutation in a cell. **B.** Hypothetical mechanism at the assembly step showing a case where mutant and wild-type proteins cannot successfully interact, preventing phenotypic mixing and allowing an increase in the number of mutant virions.

This could make adaptation at release VIPs a highly effective way to limit viral proliferation through the host, despite the higher selective constraint at release VIPs (using variants with allele frequency greater than 20%, PN/PS = 0.67 for release VIPs vs. 0.80 for VIPs on average; Figure 7). This suggests high pressure for the host to adapt, resulting in the patterns we see in release VIP genes. Additionally, we expect that we see this pattern because the virus was not stopped sooner in the viral replication cycle, perhaps because there was not a sufficient number of advantageous variants to do so. This might be more likely to happen in humans where adaptation is limited by lower effective population size and the low amount of standing genetic diversity [54–56]. This suggests that the fixed substitutions in release proteins could potentially be deleterious when not in an epidemic, because these mutations might have simply the only ones available that allowed the host to stop the virus at the very last viral replication step.

We also found the surprising result that immune VIPs do not show a significant increase in adaptation according to ABC-MK, while nonimmune VIPs do. This is contrary to many previous studies that find strong signals of positive selection at immune genes based solely on DN/DS [1, 57–59]. We hypothesize that there might be significant balancing selection happening in the immune genes studied. Balancing selection is known to be a powerful force shaping the evolution of immune genes, in particular the HLA locus [35, 60–62]. As previously mentioned, ABC-MK is expected to be biased downward in the presence of balanced nonsynonymous variants (elevated PN), thus bringing down the value of α. Due to strong prior evidence of balancing selection in a variety of immune genes, we predict that there is balancing selection occurring in the immune VIPs studied, limiting our ability to detect the presence of what is likely strong positive selection.

Because this study uses current literature as the source of our data annotations, it is necessarily limited by what host genes and viruses have been studied in the framework of molecular virology (Materials and Methods). Not only have certain viruses been studied much more extensively than others (e.g. HIV), some viral replication cycle categories have a limited number of genes that prevents more detailed analysis.

Categories like viral assembly with only 75 tested genes could possibly have significant host adaptation that would only be detectable with more genes. Future studies targeting lesser studied viruses or replication cycle categories with smaller gene counts would improve our analysis by allowing for better detection and quantification of positive selection in different categories. Additionally, further molecular research targeting the arms race between host release VIPs and viruses would help to clarify the reason for their exceptionally high observed adaptation. This adaptation that prevents new viral particles from exiting the cell to continue infection in the host, thereby trapping them in the cell, was such a frequent and important strategy in human hosts that the evidence of it can still be found in the form of extremely strong measures of positive selection.

## Materials and Methods

### Data Annotation

In order to quantify positive selection in the host at different steps of the viral replication cycle, we first annotated human VIPs using current virology literature. We began with genes encoding VIPs identified from high throughput studies. For each identified gene, we then used its associated Ensembl gene name(s) to search PubMed for articles studying the molecular function of the gene or its protein product. For each paper that characterizes the VIP, we classified the VIP as proviral (a protein that the virus uses to advance its own replication) or antiviral (a protein that the virus must suppress or evade), the step of the viral replication cycle it is associated with, and the virus it interacts with. We assigned replication cycle steps based on categories that virologists themselves have classified viral replication in the literature.

Using this approach, we obtained a list of 2,477 VIPs (43.3% of 5,724 currently known VIPs; (S1 File) with ∼7,500 proviral and antiviral replication step annotations (S1 File). This resulted in diverse functional sets of VIPs, with numbers of genes ranging from 75 to more than 1,000 (Table 1). These sets include both VIPs related to immune function and those that have no known immune function, given that immune genes have been previously found to be under strong selection [57]. Additionally, to account for the increased selective constraint on housekeeping genes, we created separate sets for VIPs that are also housekeeping genes and those that are not housekeeping genes.

The full list of replication cycle categories is as follows:

- Receptor
- **Entry** excluding receptors
- Entry and Receptor
- **Gene expression** including viral RNA transcription and translation
- Viral genome replication
- **Intermediate transport** i.e. intracellular transport after entry but before release
- **Overall Transport** including entry, intermediate transport, and release
- **Release** excluding assembly VIPs, due to the common overlap and occasional confusion between assembly and release in the virology literature
- Cell survival and/or proliferation

### Control non-VIP sets with matching confounding factors

In order to identify the adaptation that is specifically due to viruses, we used a previously described method [32, 37] to generate sets of controls consisting of non-VIPs with potentially confounding factors that match VIPs. Potentially confounding factors here are factors other than the physical interaction with viruses that can also influence the amount of protein adaptation. Using this approach allows us to compare adaptation between VIPs and non-VIPs while removing the impact of confounding factors in order to quantify only adaptation driven by viruses. The bootstrapping procedure used to generate non-VIP control sets follows the procedure thoroughly described in [32] and [37]. It uses a control set building algorithm that progressively adds control genes to a set, such that the growing control set has the same average confounding factor values for the confounding factors as the VIP set. In other words, though individual genes may not match the average for all factors, the control set as a whole matches the VIP set.

Each control set is still relatively close in values to the set of VIPs, because we set a 5% matching tolerance limit over or under each confounding factor average value. Though resampling of control genes is allowed in the algorithm, each gene is restricted to appearing 3 or fewer times in the data set.

We included, adapted from [32], the following genomic factors individually to see if they have an impact on observed positive selection enrichment across all VIPs or functional categories used in this manuscript:

- Average overall expression in 53 GTEx v7 tissues [63]. Expression in VIPs tends to be higher across many tissues than in nonVIPs [1]. To avoid confusing differential gene expression for positive selection, we match the log2 of Transcripts per Million (TPM) of the average expression.
- Average expression in GTEx lymphocytes [63]. VIPs tend to be more highly expressed in lymphocytes, thus we specifically control for differential expression between VIPs and nonVIPs again using log2 of TPM.
- Average expression in GTEx testis using log2 of TPM [63]. Genes that are highly expressed in testes tend to be enriched for signals of positive selection [64]. This is likely to impact the estimation of adaptation.
- deCode 2019 map recombination rate. Recombination rate affects the level of background selection and linkage between different new mutations. We used the most recent deCode recombination map [65] centered at the genomic center of the gene in both 50kb and 1000kb windows to account for the effect of varying window sizes (this is true for all other factors using multiple window sizes).
- GC content. GC content can be used as a proxy for biased gene conversion and recombination rate, both of which can bias estimates of adaptation [66]. We matched VIPs and non-VIPs for GC content at all three codon positions in the aligned Ensembl coding sequences [26], as well as the overall GC content.
- Proportion of human-ape orthologous CDS alignment made of indels [26].
- Number of one-to-one orthologs for a human CDS in an alignment with 535 mammals [20]. This provides an estimate of the age and level of conservation of the CDS across mammals.
- Number of one-to-one orthologs for a human CDS in an alignment with 68 primates [20]. This provides an estimate of the age and level of conservation of the CDS across primates.
- Number of protein-protein interactions (PPIs) in the human protein interaction network. VIPs tend to have a higher number of PPIs than non-VIPs, and the number of PPIs influences the rate of selective sweeps [67]. We matched the log2 number of PPIs between VIPs and control non-VIPs.
- CDS length aligned with other apes [26]. Longer coding sequences can accumulate more polymorphism and fixed substitutions because of a higher number of sites, biasing estimates of adaptation. Control non-VIP sets have the same average CDS length as the VIP sets.
- Proportion of immune genes (as annotated with GO terms ‘immune system process’, ‘defense response’, and ‘immune response’). Immune genes are well known to be hotspots of positive selection [1, 57–59], thus not controlling for the number of immune genes in each set could bias the estimate of adaptation. Control non-VIP sets have the same proportion of immune genes as the VIP sets.
- Proportion of housekeeping genes, as defined by [68]. Due to the higher selective constraint that housekeeping genes experience, adaptation in these genes is more limited, biasing estimates of adaptation when using some VIPs that are housekeeping genes and some that are not. Control non-VIP sets have the same proportion of housekeeping genes as the VIP sets.

Of all these considered possible confounders, we found that the number of protein-protein interactions, the average gene expression across tissues, gene expression in lymphocytes, the proportion of immune genes, and the number of orthologs in primates have an impact on the observed enrichment of positive selection (change in the relative selection enrichment greater than 2%). We therefore match these confounding factors between VIPs and control non-VIPs. We also match the proportion of housekeeping genes and CDS length even if we did not see an impact on the relative selection enrichment, in order to keep variance in check by making sure that control sets of non-VIPs have the same amount of sequence information as the tested sets of VIPs.

### Estimation of the False Positive Rate

We estimated the false positive rate (FPR) of our ABC-MK data by getting an approximate unbiased p-value as in [32]. We randomized the VIPs and nonVIPs for each category and performed random sampling of a set that has the same number of genes as the VIP list 1,000 times. These random sets were then run through the ABC-MK pipeline to obtain a value of α, where the number of times the random set produced a value of α higher than the VIP set was recorded. Dividing this value by the total runs (1,000) gave us an unbiased p-value for each category, which allowed us to determine the false positive rate (S1 Table). By determining the number of times the random set produced a value of α higher than the VIP set, we obtain an unbiased, nominal p-value and thus the false positive rate.

### Genomic Data Processing

Both the classical Mcdonald-Kreitman test and the ABC extension (ABC-MK) require divergence and polymorphism data to estimate the proportion of adaptive fixations in a coding sequence. The entire procedure to obtain PN, PS, DN and DS has already been described in [26]. As a brief reminder, we obtained polymorphic counts and derived allele frequencies from 1000GP phase 3 high-coverage phased data from seven African ancestry populations [69]. We then used VEP to annotate synonymous and nonsynonymous polymorphism and Ensembl v109 to estimate ancestral and derived allele frequencies from EPO multi-alignments [26]. Then we re-aligned human, chimpanzee, gorilla, and orangutan sequences with MACSE v2 to explicitly account for codon structure [70]. This CDS alignment of ape species has been previously published in [26]. From this MACSE alignment, we inferred orthology using OrthoFinder [71].

### ABC-MK

We used the MKtest Julia package [26] to apply an Approximate Bayesian Computation extension of the Mcdonald-Kreitman test to our data. ABC-MK uses α(x) statistics [25] to accurately estimate α in the presence of confounding factors such as background selection and weakly beneficial alleles. This overturns the assumption of the original MK test (and previous extensions) that beneficial mutations do not contribute to polymorphism, which can downwardly bias estimates of α when weak positive selection is pervasive. This pipeline gives an estimate of α, or the proportion of adaptive fixations in a group of genes (Equation 1) as well as separate estimates of the rate of adaptation for strongly and weakly beneficial alleles [6].

Our input model parameters to ABC-MK are as follows:

- gam_flanking (selection coefficient for deleterious alleles): −500
- gL (Selection coefficient for weakly beneficial alleles): 10
- gH (selection coefficient for strongly beneficial alleles): 500
- al_low (proportion of α due to weak selection): 0.2
- al_tot (α): 0.4
- theta_flanking (mutation rate defining BGS strength): 0.001
- theta_coding (mutation rate on coding regions): 0.001
- B (BGS strength): 0.999
- Shape (DFE shape parameter): 0. 184 [72]
- Gam_DFE: −457
- ppos_l (fixation probability of weakly beneficial alleles): 0
- ppos_h (fixation probability of strongly beneficial alleles): 0
- N (population size): 1000
- n (sample size; number of individuals): 661
- Lf (flanking region length): 2 * 10^5
- ρ (recombination rate): 0.001
- TE: 5
- Cutoff: 0.7
- DAC: [2, 4, 5, 10, 20, 50, 200, 661]

## Supporting Information

**S1 Fig.** α(x) curve for the category of gene expression

**S1 Table.** Empirical p-values from the ABC-MK pipeline shaded according to their FPR

**S1 File.** VIP functional annotations

## Acknowledgements

We thank Ryan Gutenkunst, Joanna Masel, Keith Maggert, Nathan Ellis, Todd Schlenke, and Koenraad van Doorslaer for helpful discussion.

## Author Contributions

**Conceptualization**: DE, MRW; **Data Curation**: DE, MRW, JMM**; Formal Analysis**: MRW; **Funding Acquisition**: DE; **Investigation**: MRW, DE; **Methodology**: JMM, DE; **Project Administration**: DE; **Resources**: DE; **Software**: DE, JMM; **Supervision**: DE; **Validation**: DE, MRW; **Visualization**: MRW; **Writing—Original Draft Preparation**: MRW; **Writing—Review & Editing**: MRW, DE.

## Declaration of Interests

The authors declare no competing interests.

## Notes

### Competing Interest Statement

The authors have declared no competing interest.

### Funding

- National Institute of General Medical Sciences: R35GM142677

## References

1. Enard D, Cai L, Gwennap C, Petrov DA. Viruses are a dominant driver of protein adaptation in mammals. Elife. 2016;5.

2. Harari A, Dutoit V, Cellerai C, Bart P-A, Du Pasquier RA, Pantaleo G. Functional signatures of protective antiviral T-cell immunity in human virus infections. Immunological Reviews. 2006;211(1):236–54.

3. Ortiz M, Guex N, Patin E, Martin O, Xenarios I, Ciuffi A, et al. Evolutionary trajectories of primate genes involved in HIV pathogenesis. Molecular biology and evolution. 2009;26(12):2865–75.

4. Meyerson NR, Rowley PA, Swan CH, Le DT, Wilkerson GK, Sawyer SL. Positive selection of primate genes that promote HIV-1 replication. Virology. 2014;454–455:291–8.

5. Enard D, Petrov DA. Ancient RNA virus epidemics through the lens of recent adaptation in human genomes: Ancient RNA virus epidemics. Philosophical Transactions of the Royal Society B: Biological Sciences. 2020;375(1812).

6. Uricchio LH, Petrov DA, Enard D. Exploiting selection at linked sites to infer the rate and strength of adaptation. Nat Ecol Evol. 2019;3(6):977–84.

7. Demogines A, Abraham J, Choe H, Farzan M, Sawyer SL. Dual Host-Virus Arms Races Shape an Essential Housekeeping Protein. PLOS Biology. 2013;11(5):e1001571.

8. Li W, Moore MJ, Vasilieva N, Sui J, Wong SK, Berne MA, et al. Angiotensin-converting enzyme 2 is a functional receptor for the SARS coronavirus. Nature. 2003;426(6965):450–4.

9. Wang W, Zhao H, Han G-Z. Host-Virus Arms Races Drive Elevated Adaptive Evolution in Viral Receptors. Journal of Virology. 2020;94(16).

10. Hoffmann M, Kleine-Weber H, Schroeder S, Krüger N, Herrler T, Erichsen S, et al. SARS-CoV-2 cell entry depends on ACE2 and TMPRSS2 and is blocked by a clinically proven protease inhibitor. cell. 2020;181(2):271–80.

11. Walls AC, Park Y-J, Tortorici MA, Wall A, McGuire AT, Veesler D. Structure, function, and antigenicity of the SARS-CoV-2 spike glycoprotein. Cell. 2020;181(2):281–92.

12. Asselta R, Paraboschi EM, Mantovani A, Duga S. ACE2 and TMPRSS2 variants and expression as candidates to sex and country differences in COVID-19 severity in Italy. Aging (Albany NY). 2020;12(11):10087–98.

13. Suryamohan K, Diwanji D, Stawiski EW, Gupta R, Miersch S, Liu J, et al. Human ACE2 receptor polymorphisms and altered susceptibility to SARS-CoV-2. Commun Biol. 2021;4(1):475.

14. Hussain M, Jabeen N, Raza F, Shabbir S, Baig AA, Amanullah A, et al. Structural variations in human ACE2 may influence its binding with SARS-CoV-2 spike protein. Journal of Medical Virology. 2020;92(9):1580–6.

15. Yurkovetskiy L, Wang X, Pascal KE, Tomkins-Tinch C, Nyalile TP, Wang Y, et al. Structural and Functional Analysis of the D614G SARS-CoV-2 Spike Protein Variant. Cell. 2020;183(3):739–51.e8.

16. Kuzmina A, Khalaila Y, Voloshin O, Keren-Naus A, Boehm-Cohen L, Raviv Y, et al. SARS-CoV-2 spike variants exhibit differential infectivity and neutralization resistance to convalescent or post-vaccination sera. Cell Host & Microbe. 2021;29(4):522–8.e2.

17. Zhou B, Thao TTN, Hoffmann D, Taddeo A, Ebert N, Labroussaa F, et al. SARS-CoV-2 spike D614G change enhances replication and transmission. Nature. 2021;592(7852):122–7.

18. Pontremoli C, Forni D, Cagliani R, Filippi G, De Gioia L, Pozzoli U, et al. Positive Selection Drives Evolution at the Host-Filovirus Interaction Surface. Mol Biol Evol. 2016;33(11):2836–47.

19. Diehl WE, Lin AE, Grubaugh ND, Carvalho LM, Kim K, Kyawe PP, et al. Ebola Virus Glycoprotein with Increased Infectivity Dominated the 2013&#x2013;2016 Epidemic. Cell. 2016;167(4):1088–98.e6.

20. Vazquez JM, Elise Lauterbur M, Mottaghinia S, Bucci M, Fraser D, Gaucherand L, et al. Extensive longevity and DNA virus-driven adaptation in Nearctic Myotis bats. 2025.

21. McDonald JH, Kreitman M. Adaptive protein evolution at Adh in Drosophila. Nature. 1991;351(June):652–4.

22. Kosakovsky Pond SL, Frost SDW, Muse SV. HyPhy: Hypothesis testing using phylogenies. Bioinformatics. 2005;21(5):676–9.

23. Kosakovsky Pond SL, Murrell B, Fourment M, Frost SD, Delport W, Scheffler K. A random effects branch-site model for detecting episodic diversifying selection. Mol Biol Evol. 2011;28(11):3033–43.

24. Fay JC. Weighing the evidence for adaptation at the molecular level. Trends Genet. 2011;27(9):343–9.

25. Messer PW, Petrov DA. Frequent adaptation and the McDonald-Kreitman test. Proc Natl Acad Sci U S A. 2013;110(21):8615–20.

26. Murga-Moreno J, Casillas S, Barbadilla A, Uricchio L, Enard D. An efficient and robust ABC approach to infer the rate and strength of adaptation. G3 (Bethesda). 2024;14(4).

27. Kuhl BD, Cheng V, Wainberg MA, Liang C. Tetherin and its viral antagonists. J Neuroimmune Pharmacol. 2011;6(2):188–201.

28. Neil SJ, Zang T, Bieniasz PD. Tetherin inhibits retrovirus release and is antagonized by HIV-1 Vpu. Nature. 2008;451(7177):425–30.

29. Van Damme N, Goff D, Katsura C, Jorgenson RL, Mitchell R, Johnson MC, et al. The interferon-induced protein BST-2 restricts HIV-1 release and is downregulated from the cell surface by the viral Vpu protein. Cell Host Microbe. 2008;3(4):245–52.

30. Wang X, Hinson ER, Cresswell P. The interferon-inducible protein viperin inhibits influenza virus release by perturbing lipid rafts. Cell Host Microbe. 2007;2(2):96–105.

31. Fitzgerald KA. The interferon inducible gene: Viperin. J Interferon Cytokine Res. 2011;31(1):131–5.

32. Di C, Moreno JM, Salazar-Tortosa DF, Elise Lauterbur M, Enard D. Decreased recent adaptation at human mendelian disease genes as a possible consequence of interference between advantageous and deleterious variants. eLife. 2021;10:1–27.

33. Souilmi Y, Lauterbur ME, Tobler R, Huber CD, Johar AS, Moradi SV, et al. An ancient viral epidemic involving host coronavirus interacting genes more than 20,000 years ago in East Asia. Current Biology. 2021;31(16):3504–14.e9.

34. Salvador-Martínez I, Murga-Moreno J, Nieto JC, Alsinet C, Enard D, Heyn H. Adaptation in human immune cells residing in tissues at the frontline of infections. Nature Communications. 2024;15(1):10329.

35. Ferrer-Admetlla A, Bosch E, Sikora M, Marquès-Bonet T, Ramírez-Soriano A, Muntasell A, et al. Balancing Selection Is the Main Force Shaping the Evolution of Innate Immunity Genes1. The Journal of Immunology. 2008;181(2):1315–22.

36. Zhu J, He F, Hu S, Yu J. On the nature of human housekeeping genes. Trends in Genetics. 2008;24(10):481–4.

37. Enard D, Petrov DA. Ancient RNA virus epidemics through the lens of recent adaptation in human genomes. Philos Trans R Soc Lond B Biol Sci. 2020;375(1812):20190575.

38. Li WH, Sadler LA. Low nucleotide diversity in man. Genetics. 1991;129(2):513–23.

39. Prado-Martinez J, Sudmant PH, Kidd JM, Li H, Kelley JL, Lorente-Galdos B, et al. Great ape genetic diversity and population history. Nature. 2013;499(7459):471–5.

40. Flanagan ML, Oldenburg J, Reignier T, Holt N, Hamilton GA, Martin VK, et al. New world clade B arenaviruses can use transferrin receptor 1 (TfR1)-dependent and - independent entry pathways, and glycoproteins from human pathogenic strains are associated with the use of TfR1. J Virol. 2008;82(2):938–48.

41. LeBouder F, Lina B, Rimmelzwaan GF, Riteau B. Plasminogen promotes influenza A virus replication through an annexin 2-dependent pathway in the absence of neuraminidase. J Gen Virol. 2010;91(Pt 11):2753–61.

42. Matsuyama S, Nao N, Shirato K, Kawase M, Saito S, Takayama I, et al. Enhanced isolation of SARS-CoV-2 by TMPRSS2-expressing cells. Proc Natl Acad Sci U S A. 2020;117(13):7001–3.

43. Shirato K, Matsuyama S, Ujike M, Taguchi F. Role of proteases in the release of porcine epidemic diarrhea virus from infected cells. J Virol. 2011;85(15):7872–80.

44. Ehrlich LS, Medina GN, Carter CA. Sprouty2 regulates PI(4,5)P2/Ca2+ signaling and HIV-1 Gag release. J Mol Biol. 2011;410(4):716–25.

45. Ehrlich LS, Medina GN, Carter CA. ESCRT machinery potentiates HIV-1 utilization of the PI(4,5)P(2)-PLC-IP3R-Ca(2+) signaling cascade. J Mol Biol. 2011;413(2):347–58.

46. Vazquez JM, Lauterbur ME, Mottaghinia S, Gaucherand L, Maesen S, Singer M, et al. Extensive longevity and DNA virus-driven adaptation in nearctic <em>Myotis</em> bats. bioRxiv. 2025:2024.10.10.617725.

47. Baker MR, Fan G, Serysheva, II. Structure of IP(3)R channel: high-resolution insights from cryo-EM. Curr Opin Struct Biol. 2017;46:38–47.

48. Schmitz EA, Takahashi H, Karakas E. Structural basis for activation and gating of IP(3) receptors. Nat Commun. 2022;13(1):1408.

49. Klaus JP, Eisenhauer P, Russo J, Mason AB, Do D, King B, et al. The intracellular cargo receptor ERGIC-53 is required for the production of infectious arenavirus, coronavirus, and filovirus particles. Cell Host Microbe. 2013;14(5):522–34.

50. Zeyen L, Döring T, Prange R. Hepatitis B Virus Exploits ERGIC-53 in Conjunction with COPII to Exit Cells. Cells. 2020;9(8).

51. Loverdo C, Lloyd-Smith JO. Intergenerational phenotypic mixing in viral evolution. Evolution. 2013;67(6):1815–22.

52. Charman M, Weitzman MD. Replication compartments of DNA viruses in the nucleus: location, location, location. Viruses. 2020;12(2):151.

53. Jose J, Taylor Aaron B, Kuhn Richard J. Spatial and Temporal Analysis of Alphavirus Replication and Assembly in Mammalian and Mosquito Cells. mBio. 2017;8(1):10.1128/mbio.02294-16.

54. Lynch M. Rate, molecular spectrum, and consequences of human mutation. Proc Natl Acad Sci U S A. 2010;107(3):961–8.

55. Rousselle M, Simion P, Tilak M-K, Figuet E, Nabholz B, Galtier N. Is adaptation limited by mutation? A timescale-dependent effect of genetic diversity on the adaptive substitution rate in animals. PLOS Genetics. 2020;16(4):e1008668.

56. Huber CD, Kim BY, Marsden CD, Lohmueller KE. Determining the factors driving selective effects of new nonsynonymous mutations. Proc Natl Acad Sci U S A. 2017;114(17):4465–70.

57. Shultz AJ, Sackton TB. Immune genes are hotspots of shared positive selection across birds and mammals. Elife. 2019;8.

58. Vatsiou AI, Bazin E, Gaggiotti OE. Changes in selective pressures associated with human population expansion may explain metabolic and immune related pathways enriched for signatures of positive selection. BMC Genomics. 2016;17:504.

59. Cagliani R, Forni D, Filippi G, Mozzi A, De Gioia L, Pontremoli C, et al. The mammalian complement system as an epitome of host–pathogen genetic conflicts. Molecular Ecology. 2016;25(6):1324–39.

60. Meyer D, Thomson G. How selection shapes variation of the human major histocompatibility complex: a review. Annals of Human Genetics. 2001;65(1):1–26.

61. Andrés AM, Hubisz MJ, Indap A, Torgerson DG, Degenhardt JD, Boyko AR, et al. Targets of Balancing Selection in the Human Genome. Molecular Biology and Evolution. 2009;26(12):2755–64.

62. Bitarello BD, de Filippo C, Teixeira JC, Schmidt JM, Kleinert P, Meyer D, et al. Signatures of Long-Term Balancing Selection in Human Genomes. Genome Biology and Evolution. 2018;10(3):939–55.

63. The GC, Aguet F, Anand S, Ardlie KG, Gabriel S, Getz GA, et al. The GTEx Consortium atlas of genetic regulatory effects across human tissues. Science. 2020;369(6509):1318–30.

64. Nielsen R. Molecular signatures of natural selection. Annual Review of Genetics. 2005;39:197–218.

65. Halldorsson BV, Palsson G, Stefansson OA, Jonsson H, Hardarson MT, Eggertsson HP, et al. Characterizing mutagenic effects of recombination through a sequence-level genetic map. Science. 2019;363(6425):eaau1043.

66. Duret L, Arndt PF. The impact of recombination on nucleotide substitutions in the human genome. PLoS Genet. 2008;4(5):e1000071.

67. Luisi P, Alvarez-Ponce D, Pybus M, Fares MA, Bertranpetit J, Laayouni H. Recent positive selection has acted on genes encoding proteins with more interactions within the whole human interactome. Genome Biology and Evolution. 2015;7(4):1141–54.

68. Eisenberg E, Levanon EY. Human housekeeping genes, revisited. Trends in Genetics. 2013;29(10):569–74.

69. Byrska-Bishop M, Evani US, Zhao X, Basile AO, Abel HJ, Regier AA, et al. High-coverage whole-genome sequencing of the expanded 1000 Genomes Project cohort including 602 trios. Cell. 2022;185(18):3426–40.e19.

70. Ranwez V, Douzery EJP, Cambon C, Chantret N, Delsuc F. MACSE v2: Toolkit for the Alignment of Coding Sequences Accounting for Frameshifts and Stop Codons. Molecular Biology and Evolution. 2018;35(10):2582–4.

71. Emms DM, Kelly S. OrthoFinder: phylogenetic orthology inference for comparative genomics. Genome Biol. 2019;20(1):238.

72. Boyko AR, Williamson SH, Indap AR, Degenhardt JD, Hernandez RD, Lohmueller KE, et al. Assessing the evolutionary impact of amino acid mutations in the human genome. PLoS Genet. 2008;4(5):e1000083.

