## Supplemental Figure 1 for "Stuck in the cell: trapping viruses in infected cells was a frequent adaptation strategy in human hosts"

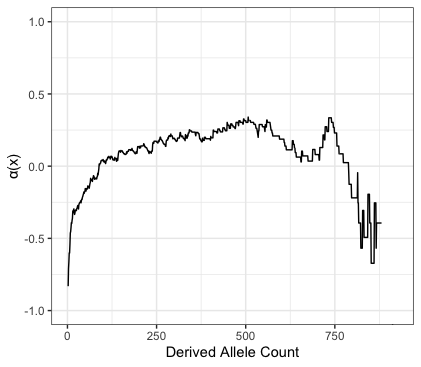

**S1 Figure.** a(x) curve for the category of gene expression. The x axis indicates the derived allele count of the 1322 total alleles in our sample population. The y axis indicates $\alpha$(x) as described in (26).
