## Supplemental Table 1 for "Stuck in the cell: trapping viruses in infected cells was a frequent adaptation strategy in human hosts"

| **Category** | **P-Value Total** | **P-Value Strong** | **P-Value Weak** |
| --- | --- | --- | --- |
| **Receptor** | **0.242** | **0.129** | **0.846** |
| **Entry** | **0.005** | **0.002** | **0.838** |
| **Entry and receptor** | **0.001** | **0.002** | **0.760** |
| **Intermediate transport** | **0.244** | **0.199** | **0.522** |
| **Replication** | **0.229** | **0.105** | **0.680** |
| **Expression** | **0.109** | **0.035** | **0.535** |
| **Transport** | **0.001** | **0.001** | **0.779** |
| **Assembly** | **0.203** | **0.258** | **0.134** |
| **Release** | **0.001** | **0.001** | **0.043** |
| **Cell Survival** | **0.309** | **0.133** | **0.859** |
| **Immune** | **0.287** | **0.561** | **0.120** |
| **Nonimmune** | **0.019** | **0.001** | **0.927** |

**S1 Table:** Empirical p-values from the ABC-MK pipeline shaded according to their FPR. Gray boxes indicate FPR < 0.05.
